# Genetic analysis of very obese children with autism spectrum disorder

**DOI:** 10.1101/185900

**Authors:** Herman D. Cortes, Rachel Wevrick

## Abstract

Autism spectrum disorder (ASD) is defined by the triad of deficits in social interactions, deficits in communication, and repetitive behaviors. Common co-morbidities in syndromic forms of ASD include intellectual disability, seizures, and obesity. We asked whether very obese children with ASD had different behavioral, physical and genetic characteristics compared to children with ASD who were not obese. We found that very obese children with ASD had significantly poorer scores on standardized behavioral tests. Very obese boys with ASD had lower full scale IQ and increased impairments with respect to stereotypies, communication and social skills. Very obese girls with ASD had increased impairments with respect to irritability and oppositional defiant behavior. We identified genetic lesions in a subset of the children with ASD and obesity and attempted to identify enriched biological pathways. Our study demonstrates the value of identifying co-morbidities in children with ASD as we move forward towards understanding the biological processes that contribute to this complex disorder and prepare to design customized treatments that target the diverse genetic lesions present in individuals with ASD.

## Introduction

Autism spectrum disorder (ASD) has complex origins with contributions from genetic and environmental factors. As more genes are identified in which sequence variants contribute to ASD predisposition, it can be useful to determine whether mutations in certain genes are responsible not only for ASD, but also for co-morbid conditions that define syndromic forms of ASD [1]. Fragile X syndrome, Angelman syndrome, Rett syndrome and a syndrome associated with *POGZ* mutations [2] are examples of syndromes with defined genetics that have high rates of behavioral phenotypes either fitting the definition of ASD or closely mimicking ASD [3]. Other syndromes with defined genetics and with elevated risks of ASD include Down syndrome [4], del16p11 syndrome [5], Bardet-Biedl syndrome [6], and Prader-Willi syndrome (PWS) [7]. However, in the latter syndromes, how the loss of specific genes confers increased risk of ASD is still unclear. For example, the expression of multiple contiguous genes is disrupted in PWS. However, mutations in only one of the inactivated genes, namely *MAGEL2*, cause a PWS-like syndrome called Schaaf-Yang syndrome [8-10]. Inactivating mutations in *MAGEL2* also carry a very high risk of ASD, suggesting that loss of *MAGEL2* also contributes to the increased incidence of ASD in children with PWS.

Children with ASD have an increased risk of overweight and obesity [11-14]. However, the genetic contribution to the co-morbidity of obesity in ASD has not been systematically studied [15, 16]. While rising rates of obesity in adults have been attributed to societal changes in physical activity and food choices, severe childhood obesity more often involves changes in the central neural circuits that regulate food intake and energy expenditure [17]. In principle, ASD and obesity could co-occur in an individual because of a single inherited or sporadic genetic mutation (e.g. *MAGEL2* deficiency), two inherited or sporadic genetic mutations (one gene responsible for ASD and a different gene responsible for obesity), or a combination of genetic mutations and multifactorial contributions. Indeed, a variety of circumstances such as reduced activity, access to health care, and interrupted sleep patterns may contribute to obesity in ASD probands [12, 14]. Some probands could have an inherited predisposition to obesity that is unrelated to their ASD diagnosis, such as variation in the melanocortin-4 receptor or FTO genes [18].

ASD- and obesity-predetermining genes participate in a large variety of biological processes, many of which affect the function of central neural circuits. Proteins important for ion transport and chromatin remodeling are common sources of mutations in autism gene sets [19]. The planar cell polarity (PCP) and Wnt-beta catenin pathways that control cell polarity and adhesion have been implicated in both autism [20-22] and in the hypothalamic regulation of energy balance [23]. A key player is the ASD-associated gene *CTNNB1*, encoding beta catenin, in which mutations cause intellectual disability (ID)/ASD with neonatal hypotonia and respiratory problems [24, 25]. Mutations in a set of genes important for PCP pathways and ciliary function cause Bardet-Biedl syndrome, a neurodevelopmental disorder associated with intellectual disability and obesity [26].

An excess of *de novo* copy number variants (dnCNVs) in probands (0.05 per individual) compared to siblings (0.02 per individual) and a larger number of genes contained within proband dnCNVs compared to sibling dnCNVs are found in autism cohorts, suggesting that dnCNVs are associated with risk of ASD [27]. Whole exome sequencing studies have estimated an average of 0.17 *de novo* likely gene disrupting (dnLGD) and 0.94 *de novo* non-synonymous coding mutations (dnNSC) per affected child, also in excess over the *de novo* events in the control siblings [27, 28]. Most recent estimates have implicated at least 8 CNVs and 65 genes with increased risk for ASD [29]. We hypothesized that candidate genes for ASD with severe obesity could be identified by analyzing genetic variants found in very obese children with ASD from the Simons Simplex Collection (SSC) [30].

## Methods

### Probands and siblings

At the time of study, the SSC Version 15 Phenotype Data Set 9 contained data on 2873 probands with ASD, their parents, and one or more siblings [31]. The data include psychological and IQ testing, medical history, dysmorphology, birth order, and anthropometric information. Details of phenotypic data are available from SFARI Base (sfari.org/resources/autism-cohorts/simons-simplex-collection). Height Z-score, body mass index (BMI), and BMI Z-score were drawn from the SSC dataset for each proband and designated unaffected sibling. As with height, normal ranges of BMI vary with age in children, so the use of Z-scores rather than absolute values allowed standardized comparison of children from 3 to 18 years of age. Height-Z and BMI-Z scores were pre-calculated in the SSC dataset using the World Health Organization growth standards and the age at evaluation from the SSC “Height, Weight, and Head Circumference” form.

### Genes

Genetic mutations in SSC probands were derived from data in GPF: Genotype and Phenotype in Families [32]. Constraint scores defining tolerance to genetic mutation were derived from Exome Aggregation Consortium (ExAC) data [33]. Intolerance to mutation is assumed when the observed number of mutations (missense or loss of function) is significantly less than the number of mutations expected.

### Statistics

Comparisons between groups of probands were performed by Student t-test followed by post-hoc comparisons with Bonferroni corrections and a value of *P*<0.05 for significance. Data in Table 1 were analyzed by Student t-test. Data in Table 2 were analyzed by the PANTHER Overrepresentation Test (release 20170413) using a Bonferroni post hoc correction for multiple testing [34]. The correlation between birth order and full scale IQ was calculated by the Pearson method. Fisher’s exact test was used to compare the mean BMI-Z and height-Z scores of probands vs unaffected siblings.

## Results

### Relationships between body mass index and anthropometric measurements, IQ, and neonatal history in SSC probands

We assessed the relationships between BMI and other anthropometric measurements, IQ, behavior, and neonatal history in the SSC SFARI Base, which contains data on probands with ASD, their parents, and siblings when available [31]. Age, height-Z, and BMI-Z scores were available for 2246 male probands and 338 female probands (Fig. 1). The mean BMI-Z of probands (0.66±1.38, mean±SD) was slightly larger than that of unaffected siblings (0.55±1.27, Student t-test, *P*=0.01). More probands than siblings had a BMI-Z≥2 (18% *vs.* 13%, Fisher’s exact test, *P*<0.0001). These data are consistent with previous reports of increased risk of obesity in children with ASD [11-14]. Mean height-Z of probands (0.4±1.1 for males and 0.2±1.2 for females) was similar to that of unaffected siblings (Student t-test).

**Figure 1.**
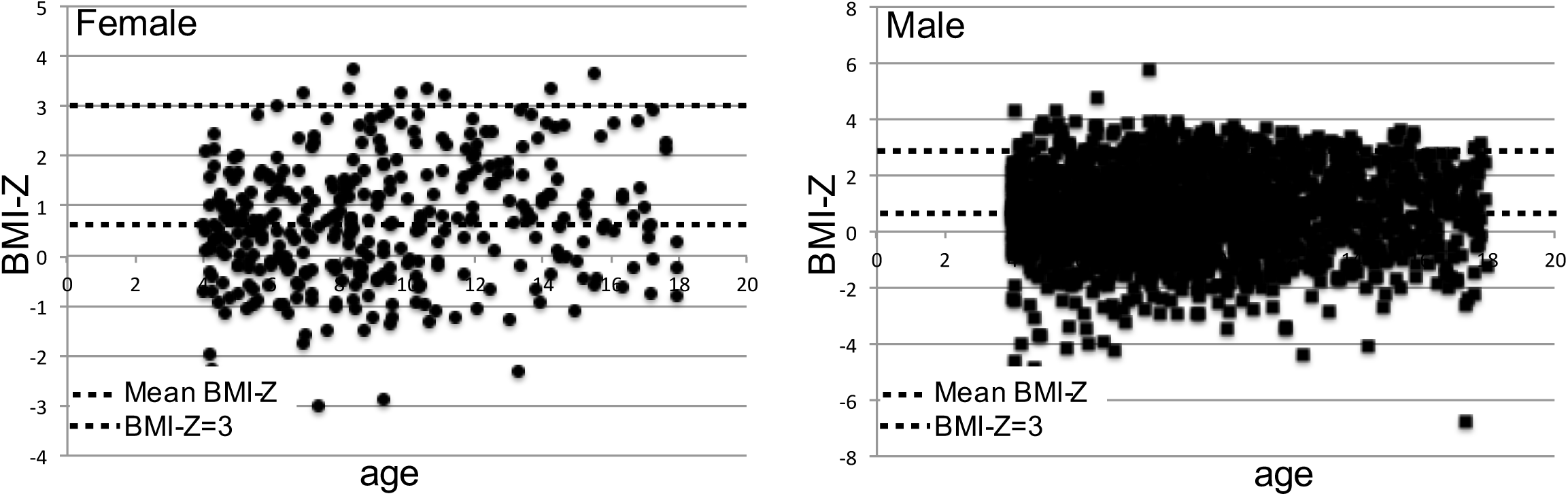
Distribution of BMI-Z versus age in the SSC.

Mother’s BMIs (mean 27±7 kg/m^2^) and father’s BMIs (mean 29±6 kg/m^2^) were calculated from their weight and height when available. Both weight and height have large genetic and environmental components, so we expected that probands with higher BMI would have parent(s) with higher BMI. Consistent with this prediction, 7% of obese probands with BMI-Z≥2 had one or two severely obese parents (BMI≥35), whereas 2.5% of probands with −2≤BMI-Z≤2 had one or two severely obese parents. This suggests that some probands with high BMI for age have an inherited predisposition to obesity, while others, particularly those with normal weight parents and siblings, may also carry deleterious mutations in obesity-predisposing genes.

We next tested whether there is a relationship between BMI-Z, ASD symptoms and full scale IQ (FSIQ) among SSC probands. In females, BMI-Z and FSIQ were not correlated. However, compared to typical weight males (-2≤BMI-Z≤2), mean FSIQ was 12 points lower among very obese males (BMI-Z≥3) (Table 1). Obese male probands (BMI-Z≥2) and underweight male probands (BMI-Z≤-2) also had significantly lower FSIQ than typical weight males. Verbal IQ was 13 points lower in very obese males compared to typical weight males. Consistent with a previous study [35], mean scores for some behaviors were worse in very obese compared to typical weight children with ASD: very obese males had increased impairments with respect to stereotypies on the Aberrant Behavior Checklist (ABC), overall language on the Autism Diagnostic Interview (ADI), social problems, somatic problems and complaints, and total competence on the Child Behavior Checklist (CBCL 6-18), awareness and communication on the teacher version of the Social Responsiveness Scale, and in communication, daily living skills, and socialization on the Vineland Adaptive Behavior Rating Scales (VABS) (Fig. 2). Very obese females with ASD had larger impairments with respect to irritability (ABC) and aggressive and oppositional defiant behaviors (CBCL) compared to typical weight females (Fig. 2). It is important to note that many of these measures are correlated with each other: for example, lower IQ is correlated with increased stereotypies [36]. There were no behaviors that were less impaired in the very obese group in any of the scales used.

**Figure 2.**
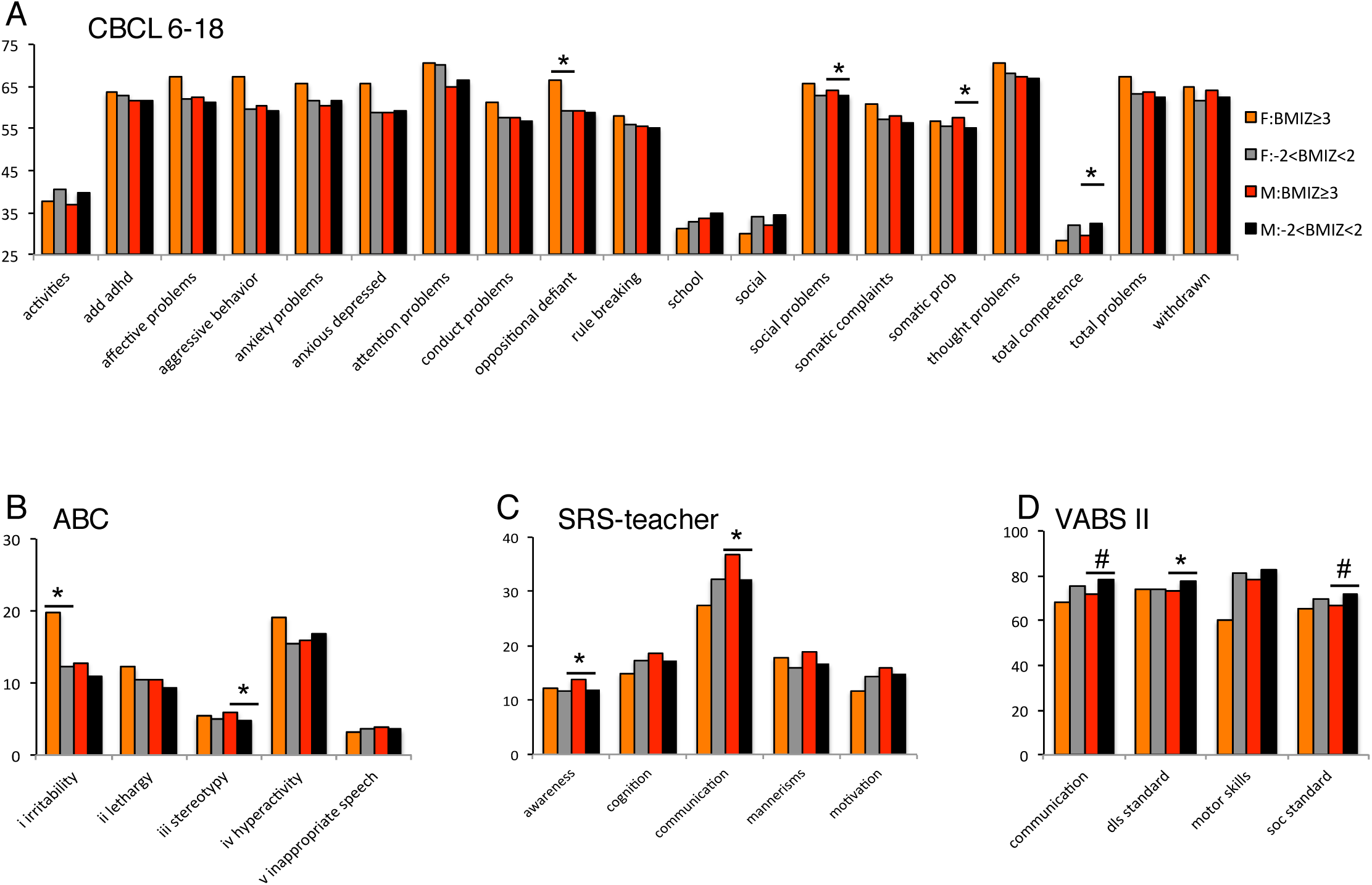
Behavioral characteristics of very obese probands on the A) Child Behavior Checklist (CBCL), B) autism behavior checklist (ABC), C) Social Responsiveness Scale (SRS), and D) Vineland Adaptive Behavior Scales (VABS). Values that are significantly different from same sex typical weight probands are indicated (\**P*≤0.01, #*P*≤0.001). Note that higher scores on the CBCL(6-18), ABC and SRS but lower scores on the VABS are indicative of behavioral deficits. Error bars are not shown for clarity.

**Table 1.** FSIQ (mean±SD) and BMI-Z in male probands.

| Group | BMI-Z | FSIQ | Verbal IQ |
| --- | --- | --- | --- |
| Group 1 (typical, n=1778) | -2<BMI-Z<2 | 83±28 | 80±31 |
| Group 2 (very obese, n=110) | BMI-Z≥3 | 72±29<sup>\$</sup> | 67±33<sup>\$</sup> |
| Group 3 (obese, n=412) | BMI-Z≥2 | 79±27<sup>*</sup> | 75±31 |
| Group 4 (underweight, n=61) | BMI-Z≤-2 | 76±28<sup>#</sup> | 73±30 |

\*P=0.002 different from Group 1
#P=0.05 different from Group 1

In humans, increased paternal age correlates with an increased number of *de novo* mutations in offspring at about 2 mutations per year [37]. However, Sanders *et al*. found no evidence for a correlation between the number of *de novo* mutations per proband and parental age in the SSC. We examined paternal age in very obese SSC probands, using birth order as a surrogate index for parental age. Very obese male probands had a higher average birth order (2.0±0.9) compared to typical weight male probands (1.7±0.8, *P*<0.0001), and so are expected to have slightly older parents. However (see below), there was no evidence that very obese probands had more *de novo* mutations than probands of typical weight.

We next examined SSC phenotypic data to determine whether obese or very obese SSC probands had other clinical findings that could suggest a syndromic form of autism. We were particularly interested in neonatal history, given that neonatal hypotonia or difficulty feeding are found in many neurodevelopmental disorders. Of 2591 SSC probands, 19% reported some neonatal complication (Fig. 3). The most common neonatal complications were difficulty feeding and respiratory problems (in 19%), poor suck (10%), or decreased muscle tone (7%). APGAR scores at 1 minute were reported to be below 7 in 5% of infants, and 7% were admitted to the NICU. These data represent minimum prevalence, as the absence of a report does not necessarily mean that the phenotype was not present. The prevalence of most neonatal complications was similar for probands who later presented as underweight (BMI-Z≤-2, n=66), obese (BMI-Z≥2, n=465), or very obese (BMI-Z≥3, n=119). There was a trend towards a higher incidence of “stiff infant” among the underweight class and a trend towards higher incidence of broken bones reported in the very obese class, but these trends were not statistically significant after accounting for multiple comparisons (Fig. 3).

**Figure 3.**
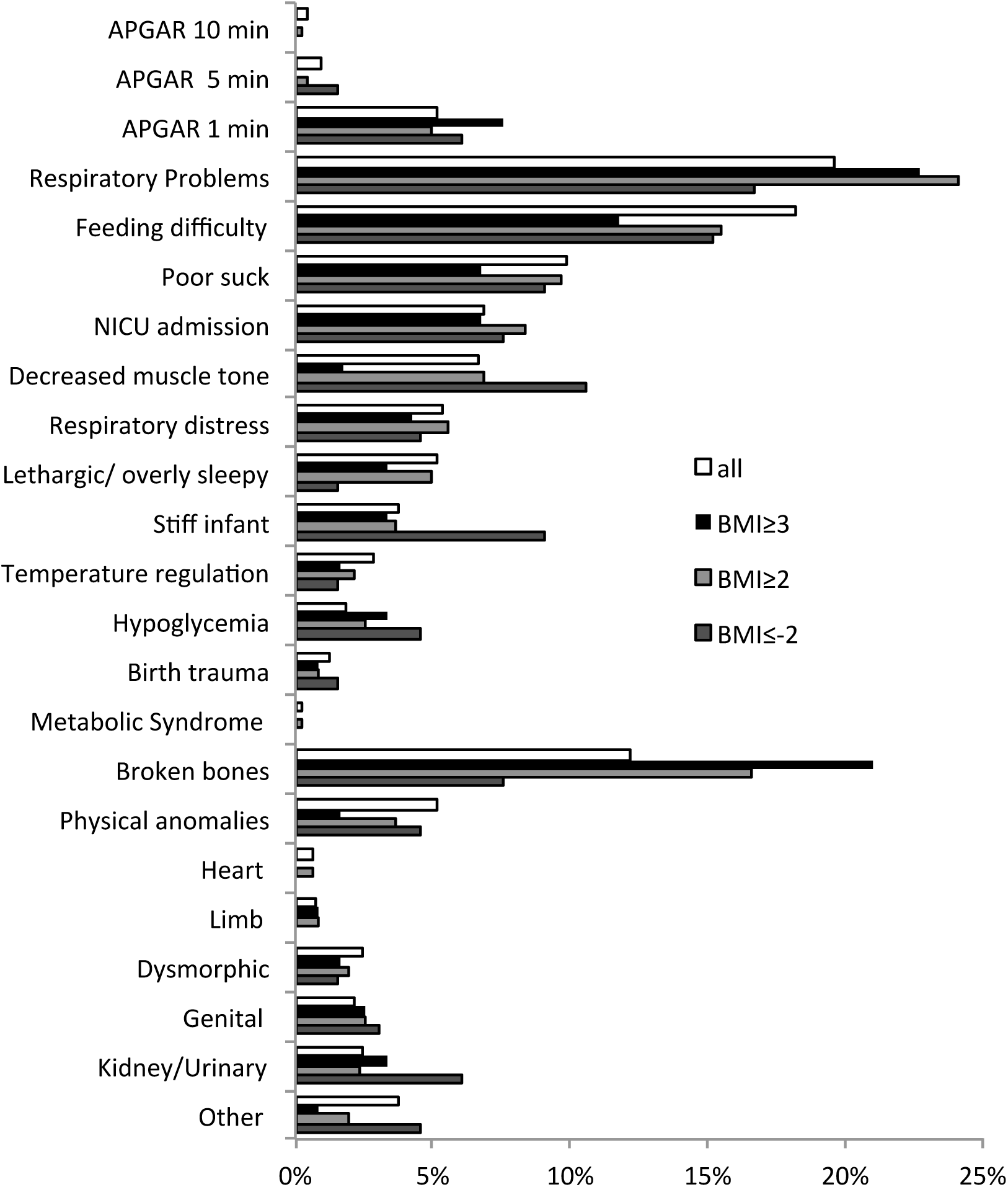
Frequency of neonatal findings in SSC probands, divided into groups by BMI-Z.

### Genetic analysis of very obese probands in the SSC

We used GPF: Genotype and Phenotype in Families to identify *de novo* mutations in 133 probands with BMI-Z≥3 (10 F, 123 M) [32]. We limited the search to variant types sub, ins, del, or CNV with effect types coding (Nonsense, Frame-shift, Splice-site, Missense, Non-frame-shift, noStart, noEnd) or CNV (+ or -). There were 120 such *de novo* mutations, including 17 dnCNVs, 20 dnLGD, Frame-shift, Nonsense, or Splice-site) mutations and 83 mutations of the remaining varieties (Suppl. Table 1). The number of *de novo* mutations per very obese proband was not different from the number of *de novo* mutations per typical weight proband. Some probands had *de novo* mutations in more than one category. Three dnCNVs in very obese probands had already been associated with ASD risk [38-41] (dup7q11, dup15q12, del16p11). Both the dup15q12 and del16p11 CNVs are also associated with obesity [42].

#### Analysis of loss of function mutations

Of the 20 genes with dnLGD mutations in very obese probands, 11 had already been associated with ASD in SFARI Gene [31] (*ANK2, DNMT3A, DSCAML1, MYH10, POGZ, PPM1D, PSD3, PTK7*), OMIM (*VCP*) or in neurodevelopmental disorders [2, 43] (*SPEN*, *STAM*). Only two of these genes have been associated with overgrowth: *DNMT3A* and *POGZ*. The proband identified here is the only one in the SSC with a dnLGD mutation in *DNMT3A* (DNA-methyltransferase 3 alpha). *DNMT3A* is mutated in children with Tatton-Brown-Rahman syndrome, which is associated with tall stature, a distinctive facial appearance, and intellectual disability. *De novo* inactivating mutations in *POGZ*, encoding the chromatin factor Pogo transposable element with ZNF domain, cause syndromic autism with hypotonia and obesity (White-Sutton syndrome [2]). Only two probands in the SSC have dnLGD mutations in POGZ: the one identified here with BMI-Z=3.6 and another with BMI-Z=2, both consistent with White-Sutton syndrome. Inactivating mutations in *PTK7* (encoding a tyrosine-protein kinase), and inherited or *de novo* mutations in *PPM1D* (encoding a protein phosphatase) have been found in children with ASD [27, 44]. The female with a *PPM1D* mutation also carries a dnNSC in *SCN1A*, encoding a sodium channel associated with epilepsy, and a dnLGD mutation in *TNFRSF8*. dnLGD mutations in *MYH10* (encoding a non-muscle myosin) have been found in children with moderate to severe ID [44-47]. Heterozygous LGD mutations in *VCP* (valosin containing protein) cause amyotrophic lateral sclerosis with dementia or myopathy [48]. The male with a *VCP* dnLGD mutation also carried a dnNSC in *SCN4A*, a gene mutated in autosomal dominant and autosomal recessive muscular conditions. The other 10 genes in which dnLGD mutations were found have not been associated with neurodevelopmental genetic disorders to date. There were 83 dnNSC mutations, of which 20 are in genes plausibly associated with autism in SFARI gene (*ANK3*, *CSMD1, DISC1, GLRA2, GRM7, KMT2C, MCC, NLGN2, PRKDC, PTK7, SCN1A, SCN4A, SLCO1B3, TRIP12, TTN, TUBGCP5, ZC3H4)* or other neurodevelopmental disorders *(ANO3, CNTN6, C9orf156, RYR1, SCN11A, SPEN, TAF6, TRPM6*). Variants in *ZC3H4* are also associated with uncontrolled eating [49], but otherwise none of the genes with dnNSC are associated with obesity.

#### Consideration of tolerance to mutation for ASD/obesity candidate genes

We next considered the tolerance of the candidate genes to either loss of function (LoF) or missense mutations using the ExAC interface [33]. Reduced tolerance would be expected of a gene for which haploinsufficiency reduces fertility, and would increase the likelihood that the *de novo* mutation is pathogenic. Of the genes with dnLGD mutations, ten previously identified ASD/ID genes were intolerant to LoF, missense or both (*ANK2, DNMT3A*, *DSCAML1, MYH10, POGZ, PPM1D*, *PTK7*, *SPEN*, *STAM*, and *VCP*). Of the remaining dnLGD genes, some were highly intolerant to LoF mutations (*CDC23, SPAG9*) or to missense mutations (*CDC23*) while the remaining were tolerant to both LoF and missense mutations. Of the genes in which there were dnNSC mutations in very obese probands, some were intolerant to either missense or LoF mutations, including genes associated with neurodevelopmental disorders (*GLRA2*, *GRM7*, *KMT2C*, *MEGF8, PTK7*, *RYR1, SCN1A, TRIP12*). Other genes not yet associated with disorders were also deemed intolerant to either missense or LoF mutations, while the remaining genes were either tolerant or of unknown tolerance (Suppl. Table 1).

#### Gene ontology pathways identifies proteins important for neuronal action potentials as contributors to ASD in very obese probands

We used Gene Ontology (GO) to identify pathways enriched in the *de novo* mutated gene set compared to expected for all GO-annotated genes in the genome [50]. This analysis is constrained by the fact that the very obese group differs from the typical weight group not only in high BMI-Z but also in lower IQ (Table 1) and more severe behavioral scores (Fig. 2), so that the genes mutated in this group may contribute to ASD or any one of these co-morbid phenotypes. The combined set of mutated genes (dnLGD and dnNSC) is significantly enriched in genes involved in neuronal membrane depolarization during action potential and voltage-gated ion channel activity (*ANK2, ANK3, ANO9, ELK1, GLRA2, GRM7, NLGN2, RYR1, SCN1A, SCN3A, SCN4A, SCN11A, SLC17A3, TRPM6, VCP*) (Table 2). Analysis of missense mutations alone produced a similar profile of disruption of neuronal action potential.

**Table 2.** Gene Ontology Consortium Enrichment Analysis.

| GO biological process complete | Fold Enrichment | P value |
| --- | --- | --- |
| 1: neuronal action potential | 36 | 0.0003 |
| 1a: regulation of membrane potential | 6 | 0.003 |
| 2: membrane depolarization during action potential | 25 | 0.01 |
| GO molecular function complete |  |  |
| 3: voltage-gated sodium channel activity | 36 | 0.02 |
| 3a: sodium channel activity | 30 | 0.004 |
| 3b: voltage-gated ion channel activity involved in regulation of postsynaptic membrane potential | 58 | 0.003 |
| 3bi: gated channel activity | 8 | 0.02 |

Data were derived from PANTHER Overrepresentation Test (release 20170413), using the GO Ontology database released 2017-06-29, and a Bonferroni correction for multiple testing.

#### Planar cell polarity / Wnt signaling genes are also mutated in obese probands

We next investigated whether any of the mutated genes were previously implicated in obesity pathways. Intriguingly, all three SSC probands who carry *de novo* mutations in the PCP/Wnt pathway *PTK7* gene [51] are very obese (Suppl. Table 2). PCP pathways intersect with phosphoinositide 3-kinase (PI3K) pathways, and 3 additional very obese probands carry *de novo* mutations in the PI3K pathway genes *PSD3, MYH10*, and *ANK3* respectively. We further investigated PCP/Wnt pathway genes in obese probands in the SSC (i.e. BMI-Z≥2). An obese SSC proband (BMI-Z=2.5) carries a dnLGD mutation in *DVL3*, one of the three human genes encoding forms of the Wnt relay protein Dishevelled [52]. This proband had neonatal feeding difficulties and physical anomalies including genital anomalies. Another obese SSC proband (BMI-Z=2.5) carries a dnLGD mutation in *NOTCH1*, encoding a Dishevelled antagonist. This proband had neonatal feeding difficulties, and was stiff at birth, phenotypes also seen in infants with Adams-Oliver syndrome caused by *NOTCH1* mutations [53]. *CNTN6* encodes a NOTCH signaling pathway protein (contactin) and is mutated in a very obese proband [54]. Yet another obese SSC proband (BMI-Z=2.7) carries a dnLGD mutation in *MED13L*, a gene associated with ID with transposition of the great arteries, and implicated in Wnt signaling [55]. This proband had neonatal respiratory distress, feeding difficulty, and other unspecified defects. Last, we looked at the *DYRK1A* gene, encoding a kinase that inactivates GSK3-beta and is implicated in neurodevelopmental disorders with co-morbid autism [56]. Of four probands in the SSC with *DYRK1A* mutations, three were obese and two had multiple neonatal complications. The Wnt/PCP genes *PTK7, RAPGEF2, CTNNB1, DVL3, MED13L, DYRK1A*, and *NOTCH1* are all highly intolerant to LoF mutations, supporting their essential role in development. There were no loss of function mutations in the SSC collection in genes responsible for Bardet-Biedl syndrome, which is also a disorder of planar cell polarity. This is likely because children with Bardet-Biedl syndrome typically develop blindness and deafness, which would exclude them from the SSC.

We next considered a potential role for the *MUC* genes in ASD with high BMI. The encoded mucin proteins are aberrantly regulated through Wnt-dependent pathways in tumors, particularly in cancers of the gastrointestinal tract. Variants in MUC5B are associated with type 2 diabetes [57]. Recurrent *de novo* nonsynonymous mutations in the *MUC5B* gene have previously been noted in cohorts of patients with autism or intellectual disability. However, *MUC5B* has been previously excluded as an ASD candidate gene because of high variation in controls [58]. In contrast, ExAC analysis suggests that *MUC5B* is highly constrained for both missense and LoF mutations despite a high absolute number of variants in controls. All five probands in the SSC collection carrying *MUC5B* mutations had neonatal complications, and two were obese with BMI-Z=3.0 (missense mutation) and BMI-Z=2.9 (nonsense mutation) (Supplementary Table 3).

Last, genes encoding proteins important for endosomal sorting, protein ubiquitination and autophagy, pathways recently implicated in neurodevelopmental and neurodegenerative disorders [59] were also analyzed further. This analysis was motivated by the finding that mutations in the *MAGEL2* gene cause ASD in Schaaf-Yang syndrome and contribute to obesity and ASD in Prader-Willi syndrome through disruption of endosomal protein trafficking [8-10, 60]. Genes associated with these pathways and mutated in very obese probands include *VCP* (valosin containing protein, mutations associated with ubiquitinated inclusions and protein aggregates in cells, frontotemporal dementia with myopathy, vesicle transport and fusion), *STAM* (component of the endosomal sorting complex), and *SPAG9/JIP4* (kinesin-mediated transport in neurons).

## Discussion

The Simons Simplex collection is a well-characterized cohort of individuals with autism spectrum disorder, for whom a rich variety of clinical data has also been collected. By its nature, individuals with ASD but also a diagnosis of a syndromic genetic disorder, such as Prader-Willi syndrome or Bardet-Biedl syndrome, are underrepresented in or excluded from SSC. As observed in other autism cohorts, probands in the SSC are overweight compared to unaffected siblings. We postulated that probands in the SSC at extreme ends of the phenotypic ranges for various traits could inform us about as yet unrecognized subtypes of ASD. Indeed, we found that very obese male SSC probands had lower full scale and verbal IQ compared to typical weight probands, and also had significantly worse scores in the areas of communication, daily living skills, and socialization. More severe behaviors were also seen in very obese female probands, but the smaller number of girls compared to boys represented in the SSC limited this analysis. We cannot exclude the possibility that being very obese in and of itself impairs social communication because of exclusion from activities with peers. Conversely, more severe behaviors could contribute to obesity because the child may be excluded from activities that promote fitness. Our study is also compromised by the considerable environmental contribution to the obesity phenotype, particularly in a country that has seen childhood obesity rates increase substantially in the last few decades. Limiting the analysis to very obese probands with two lean parents might have mitigated some of this confounding variable, but would have also reduced the sample size to an unmanageably small number of probands. We did not find that neonatal complications were associated with obesity in this study, although the lack of reporting of neonatal histories limited this analysis.

We proposed that genetic investigation of individuals with dual diagnoses of ASD and obesity could point us towards pathways that are essential for normal neurodevelopment and normal development of energy-regulating pathways. Our focus on *de novo* mutations in very obese probands correctly identified obesity predisposing lesions such as del16p11 and loss of function mutations in *DNMT3A* and *POGZ* and support a role for mutations in *PTK7* in autism [40] and also in obesity. Additional very obese probands carried mutations in genes implicated in ASD or other neurological disorders, such as those important for neuronal action potentials, as predicted from previous autism analyses [61, 62]. Obese probands do not appear to have more pathogenic mutations compared to typical weight probands with ASD. Despite analyzing over 2000 probands, we could not determine whether some probands had mutations in two different genes or environmental factors that independently contributed to ASD and obesity. We also only examined *de novo* mutations, and so do not have information about other genetic mechanisms, such as recessive mutations that may contribute to either the obesity or ASD phenotypes.

The PCP/Wnt pathway has been implicated in neurodevelopmental disorders associated with neonatal complications and obesity, in part through the identification of key genes such as *CTNNB1, MIB1, RAB23, MEGF8, PTK7*, and the *BBS* genes in syndromic forms of intellectual disability*. PTK7* encodes protein tyrosine kinase 7 and is implicated in Wnt signaling pathway. PTK7 (CCK4) binds intracellular scaffolding proteins of the Dishevelled family that act downstream of WNT receptors. PTK7 has mostly been studied in the context of cancer cell motility and in epithelial to mesenchymal transitions during development. A recessive *Ptk7* mutation in mice (*chuzhoi*) causes severe developmental defects consistent with dysregulation of PCP, but no phenotype has been described in *chuzhoi* heterozygotes. Loss of *ptk7* function in zebrafish causes ciliary and PCP abnormalities. In humans, we would not expect to find homozygous mutations, considering that this state is embryonic lethal in mice. It would however be interesting to determine whether *chuzhoi* heterozygous mice have behavioral abnormalities recapitulating ASD. The endosomal sorting/ubiquitination pathways have also been implicated in syndromic intellectual disability, with mutations in *MAGEL2, VPS13B*, and *VCP* causing syndromic neurological disorders. Our data support the previously noted involvement of both the PCP/Wnt pathway and the endosomal sorting /ubiquitination pathway in ASD. There are too few probands with obesity and ASD and *de novo* mutations to determine whether this pathway is indeed enriched in the very obese subgroup of children with ASD. Further investigation of these two pathways in ASD may be warranted. In particular, genes that are mutated in obese probands with ASD and in many cases also intolerant to mutation (*BTBD9, DVL3, MED13L, RAPGEF2, RYK, STAM*, and *SPAG9*) may be good candidates for involvement in syndromic forms of ASD.

## Conclusions

Our study provides a model for gene discovery using additional phenotypic information to categorize subsets of ASD probands. For example, probands with dysmorphic features could have a different set of genetic lesions compared to those without dysmorphic features. Extensive phenotypic characterization and collection of a larger cohort of children with ASD, as is now ongoing in the SPARK initiative [63] will enrich our understanding of the multitude of genes associated with the many flavors of autism spectrum disorder. This understanding is essential for the design of customized therapies that target the underlying genetic lesion that contributes to the diagnosis of ASD in each individual.

## List of Abbreviations

(ABC): Aberrant Behavior Checklist
(ADI): Autism Diagnostic Interview
(ASD): Autism spectrum disorder
(BMI),: Body mass index
(CBCL 6-18): Child Behavior Checklist
(dnCNVs): De novo copy number variants
(dnLGD): De novo likely gene disrupting
(dnNSC): De novo non-synonymous coding mutations
(ExAC): Exome Aggregation Consortium
(FSIQ): Full scale IQ
(GO): Gene Ontology
(ID)/: Intellectual disability
(LoF): Loss of function
(PCP): Planar cell polarity
(PWS): Prader-Willi syndrome
(SSC): Simons Simplex Collection
(VABS): Vineland Adaptive Behavior Rating Scales

## Declarations

### Ethics approval and consent to participate

Use of SSC data from human subjects was approved by the Health Research Ethics Board-Biomedical Panel of the University of Alberta, Edmonton, Alberta Canada.

### Availability of data and material

The data that support the findings of this study are available from the Simons Foundation but restrictions apply to the availability of these data, which were used under license for the current study. Data are however available from the authors upon reasonable request and with permission of the Simons Foundation.

### Authors’ contributions

HDC and RW analyzed and interpreted the mutation data and wrote the manuscript. Both authors read and approved the final manuscript.

## Acknowledgements

We are grateful to all of the families at the participating Simons Simplex Collection (SSC) sites, as well as the principal investigators (A. Beaudet, R. Bernier, J. Constantino, E. Cook, E. Fombonne, D. Geschwind, R. Goin-Kochel, E. Hanson, D. Grice, A. Klin, D. Ledbetter, C. Lord, C. Martin, D. Martin, R. Maxim, J. Miles, O. Ousley, K. Pelphrey, B. Peterson, J. Piggot, C. Saulnier, M. State, W. Stone, J. Sutcliffe, C. Walsh, Z. Warren, E. Wijsman). We appreciate obtaining access to phenotypic data on SFARI Base. Approved researchers can obtain the SSC population dataset described in this study by applying at https://base.sfari.org. We are grateful to the Prader-Willi Association of Alberta for their donation in support of our research on autism spectrum disorders.

---

**Supplementary Table 1.**
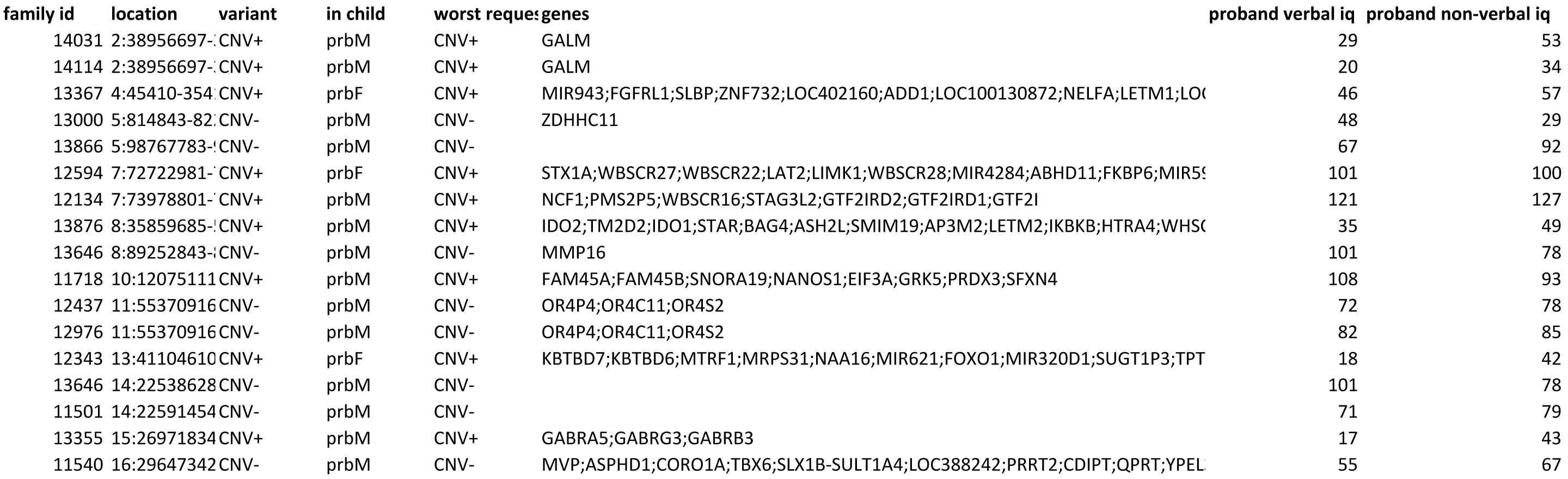
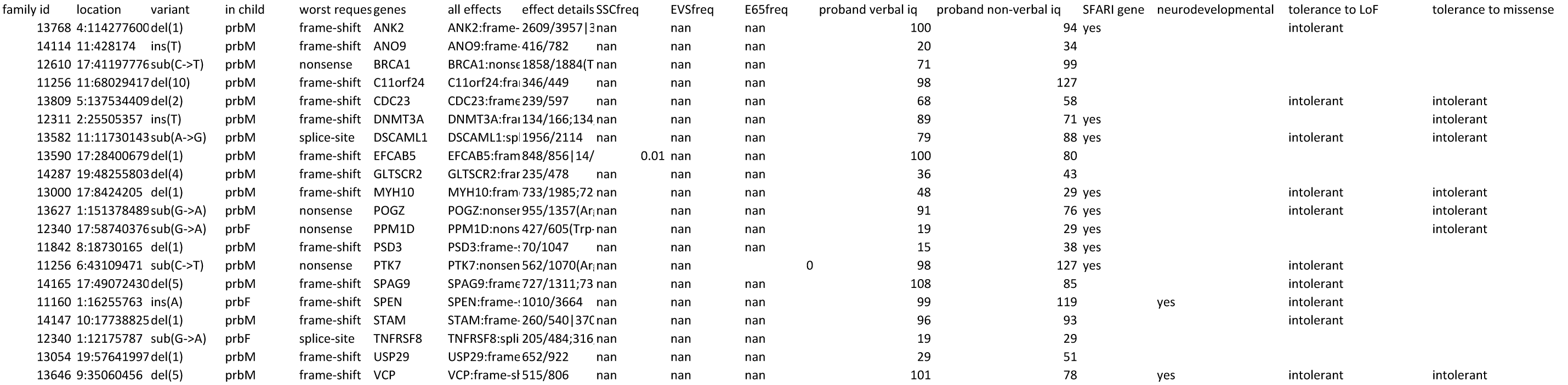
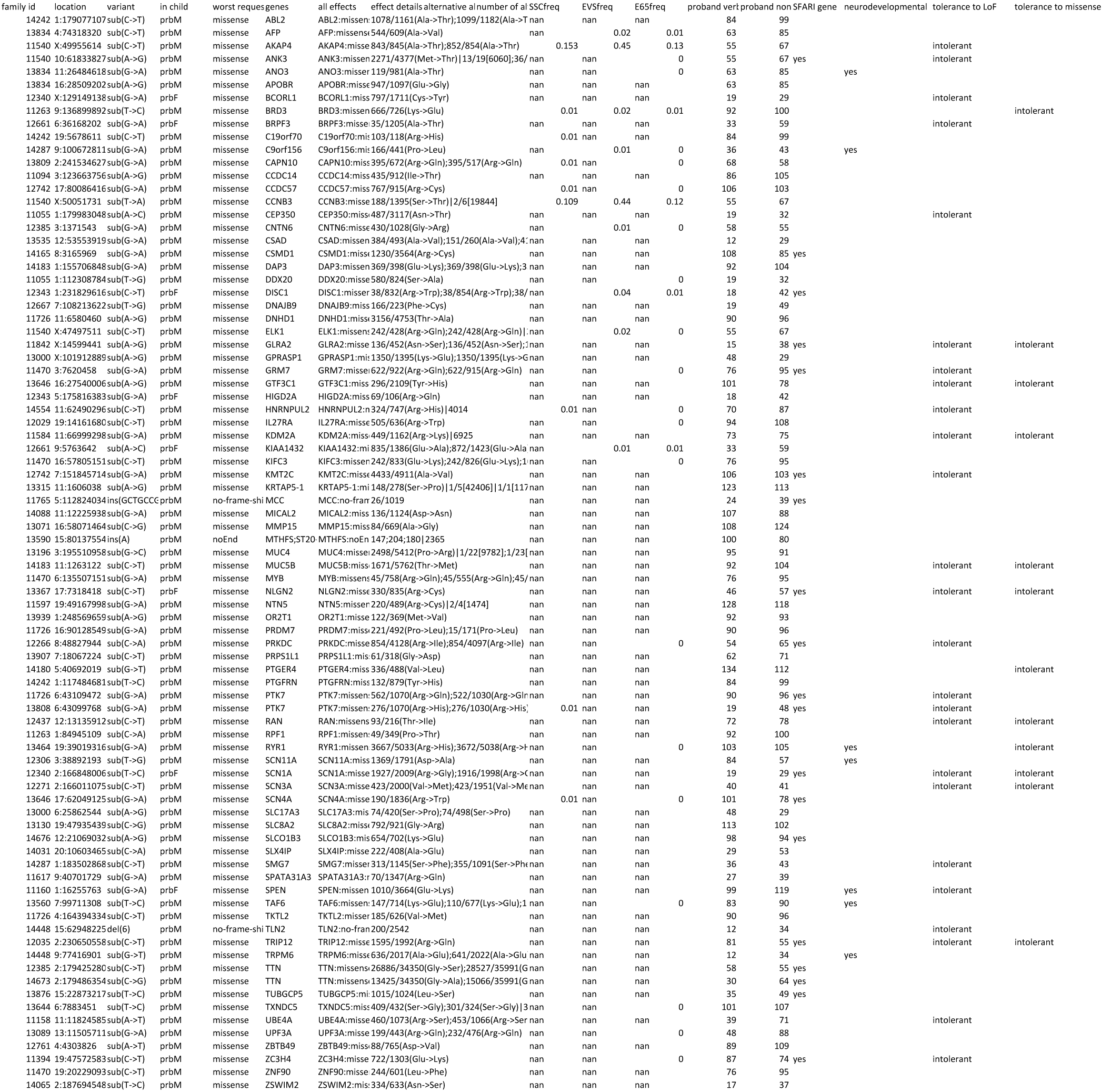
*De novo* genetic events detected in very obese probands (BMI-Z≥3). Supplementary Table 2. Phenotypes associated with *de novo* mutations in *PTK7, MEGF8* and *DVL3*. Supplementary Table 3. Phenotypes associated with *de novo MUC5B* mutations.

**Supplementary Table 2.**
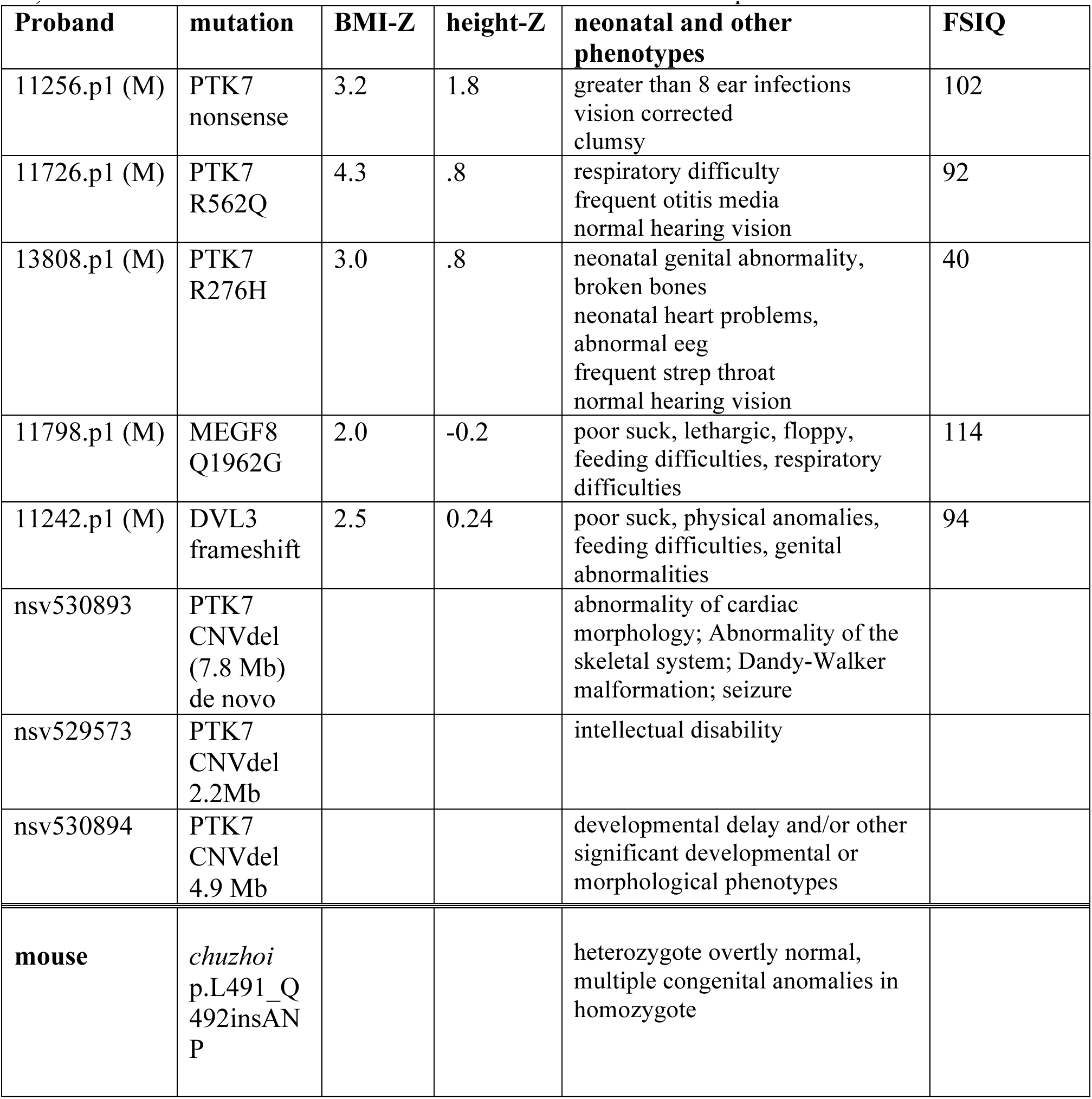
Mutations in *PTK7, MEGF8* and *DVL3* in the SSC (rows 1-5, proband IDs are masked) or the Database of Genomic Variants (rows 6-8, Kamisnsky et al., Genet Med. 2011:777-84.). The mouse mutant chuzhoi is listed in the bottom row for comparison.

**Supplementary Table 3.** Phenotypes associated with *de novo MUC5B* mutations in the Simons Simplex Collection.

| **Proband (sex)** | **Mutation** | **BMI-Z** | **height-Z** | **neonatal phenotype** | **FSIQ** |
| --- | --- | --- | --- | --- | --- |
| 11078.p1 (M) | nonsense | 2.93 | .99 | stiff infant lethargic/overly<br>sleepy large head circ | 29 |
| 14183.p1 (M) | T1671M | 3.01 | 3.47 | feeding<br>clumsy<br>broken bones<br>>8 otitis | 100 |
| 12779.p1 (M) | P4997L | 1.91 | 1.15 | feeding<br>vision corrected | 99 |
| 13983.p1 (F) | P3887T | -1.75 | .28 | feeding | 40 |
| 11653.p1 (M) | del(15) no<br>frame shift<br>V2574A<br>T3883M | -.43 | -.77 | stiff infant | 36 |
| 11247.p1 (M) | P4997L | -0.6 | +0.4 | kidney or urinary | 128 |
| 11957.p1 (M) | C4174F | 1.6 | -2.0 | physical anomalies, respiratory | 77 |
| 12447.p1 (M) | H4050Q | 0.4 | -0.5 | lethargic | 53 |

